# Panic at the Discordance: the Asparagaceae Phylogenetic Backbone

**DOI:** 10.64898/2026.09.30.755722

**Authors:** Philip C. Bentz, Ethan Baldwin, Mario Martínez-Azorín, Kálmán Könyves, Timothy S. Harvey, Joanne Birch, Etelvina Gándara, Bryan MacNeill, Andrew Salywon, Xiaohan Yang, Sierra Jaeger, James H. Leebens-Mack, Cody Coyotee Howard

**Author notes:** Corresponding author: Cody Coyotee Howard, 4110 Libra Dr., Orlando, FL 32816.

## Abstract

The Asparagaceae is an economically and ecologically important angiosperm (monocot) clade, containing iconic garden plants such as asparagus, hyacinth and agave, and arid-adapted keystone species like Joshua tree and dragon blood tree. The approximately 2,500 species in the family are spread across seven subfamilies: Agavoideae, Aphyllanthoideae, Asparagoideae, Brodiaeoideae, Lomandroideae, Nolinoideae and Scilloideae. Relationships among the seven subfamilies are generally unresolved and the sources of gene tree-species tree discordance are largely unknown. Investigating the biology underlying such discordance requires a robust sampling of both taxa and molecular markers, due to the extremely short branches that form the backbone of the Asparagaceae phylogeny. Using targeted sequencing of 1,726 low-copy nuclear genes, and a nearly complete sampling of genera, we interrogate relationships among all seven Asparagaceae subfamilies and investigate the underlying sources of discordance across the phylogeny. Analyses using different phylogenomic approaches (multispecies coalescence vs. nuclear gene supermatrix vs. chloroplast gene supermatrix) each revealed extensive discordance across the Asparagaceae backbone, which may be associated with rapid lineage diversifications that occurred in Africa shortly following the origin of the family ∼49 Ma. Specifically, we found that two branches, representing ancestors of Aphyllanthoideae and Nolinoideae, may be responsible for much of the backbone discordance. The placement of Aphyllanthoideae, represented by one extant taxon (*Aphyllanthes monspeliensis),* is controversial with discordant topologies across many studies. However, multispecies coalescence-based analyses from this study strongly support a sister relationship between Aphyllanthoideae and a larger Agavoideae + Scilloideae + Brodiaeoideae clade. According to gene tree quartet frequencies and ABBA-BABA tests, we cannot reject incomplete lineage sorting (ILS) as the main source of discordance for the placement of Aphyllanthoideae; our results leave open the possibility of an ancient reticulation involving an ancestor of Aphyllanthoideae and an extinct basal Asparagaceae lineage. Regarding Nolinoideae, we found that conflicting support for two possible bifurcating clades (Asparagoideae + Nolinoideae vs. Asparagoideae + Lomandroideae) was associated with skewed support for alternative gene tree topologies, suggesting that phenomena other than ILS may explain the observed discordance. A quartet-based ABBA-BABA analysis, used to test for signatures of introgression, revealed a putative ancient hybridization event between two divergent, ancestral lineages which may have contributed to the origin of the Nolinoideae clade in Africa ∼45 Ma. In sum, this study elucidates the long-troublesome backbone of the Asparagaceae phylogeny and advances our understanding of the evolutionary and biogeographical origins of all extant subfamilies. More broadly, this study expands our understanding of the various sources of gene tree discordance that may arise from natural processes associated with rapid lineage radiations across the tree of life.

## INTRODUCTION

A myriad of factors confound our understanding of the evolutionary relationships across the tree of life. From deep-time to more recent timescales, various biological processes can lead to discordance between the true species phylogeny and individual gene trees, with hybridization, lineage-specific de novo evolution, incomplete lineage sorting (ILS), and whole genome duplications (WGDs) as significant contributors (Pease et al. 2016; McLay et al. 2023). These processes can both uniquely and jointly obscure phylogenetic relationships, but nonetheless contribute to lineage and morphological diversification (Landis et al. 2018; Han et al. 2020; Parins-Fukuchi et al. 2021; Stull et al. 2021, 2023; Feng et al. 2024). The angiosperm family Asparagaceae (monocots) consists of approximately 2,500 species across seven taxonomic subfamilies: Agavoideae, Aphyllanthoideae, Asparagoideae, Brodiaeoideae, Lomandroideae, Nolinoideae, Scilloideae (Chase et al. 2009; Chase 2016). Species richness across these seven subfamilies varies drastically, with Scilloideae having the highest at 1,000+ accepted taxa, and Aphyllanthoideae being monotypic with only one extant species—*Aphyllanthes monspeliensis* L. (Buerki et al. 2012; Seberg et al. 2012). Establishing relationships at both deep and shallow phylogenetic scales has sometimes been fraught with uncertainty due to the aforementioned biological processes (McKain et al. 2012; Seberg et al. 2012; Givnish et al. 2018; Gunn et al. 2020, 2024; Meng et al. 2020; Heyduk et al. 2021; Howard et al. 2023; Bentz et al. 2024; Hall et al. 2026). Although previous studies have improved phylogenetic resolution among taxa within subfamilies, relationships among the subfamilies and some genera generally remain unresolved (Fig. 1). Investigating gene tree discordance across the phylogenetic backbone of Asparagaceae requires a robust sampling of both taxa and molecular markers (coalescent units or independently segregating loci). In doing so, the results will advance our understanding of evolutionary processes contributing to adaptive variation that may arise during rapid lineage radiations.

**Figure 1.**
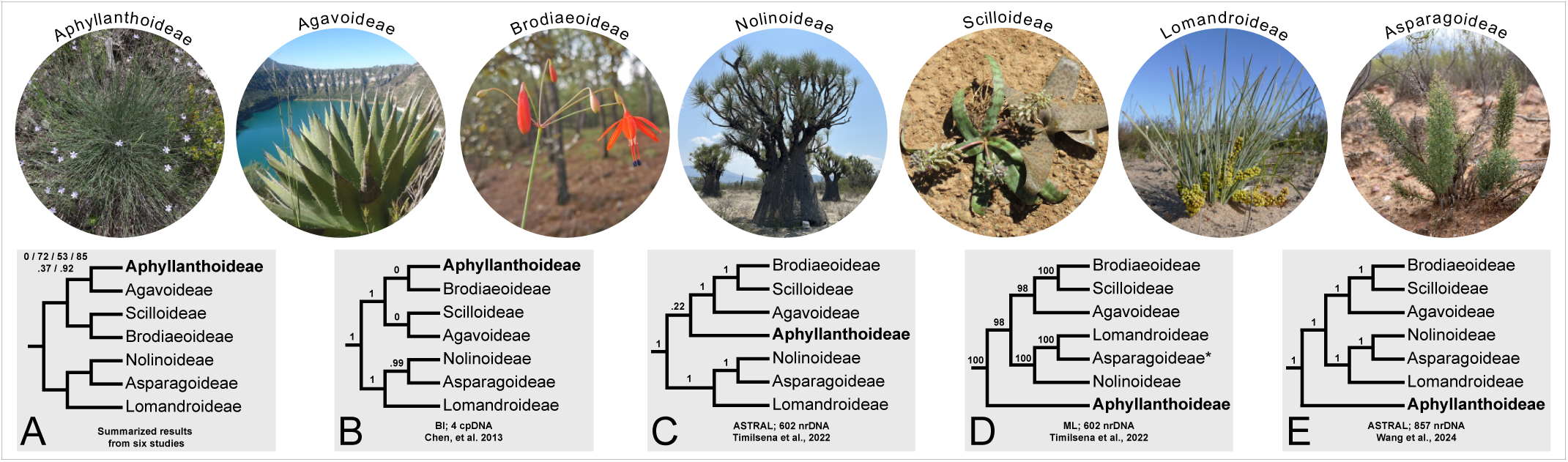
Aphyllanthoideae (bolded) changes phylogenetic placement across studies. A) Six separate studies reconstructed Aphyllanthoideae sister to Agavoideae. Support values are in the upper left corner and indicate support for the placement of Aphyllanthoideae. Maximum Likelihood (ML) bootstrap support values (top row) from Seberg, et al. 2012, (Steele et al. 2012), Givnish, et al. 2018, Ji, et al. 2023; BI local posterior probability (bottom row) from (Kim et al. 2010) and Ji, et al. 2023. B) Aphyllanthoideae placement using four chloroplast loci and bayesian inference (BI) methods from Chen, et al. 2013. Support values are local posterior probability. C) Aphyllanthoideae placement using 602 nuclear loci and species-tree methods (ASTRAL) from Timilsena et al., 2022. Support values are local posterior probability. D) Aphyllanthoideae placement using 602 nuclear loci and ML phylogenetic tree reconstruction methods from Timilsena et al. 2022. Support values are parametric bootstrap values. E) Aphyllanthoideae placement using 857 nuclear loci and species-tree methods (ASTRAL) from (Wang et al. 2024). Support values are local posterior probability. The representative taxon per subfamily is: Aphyllanthoideae—*Aphyllanthes monspeliensis* L. (photo credit: Mario Martínez-Azorín), Agavoideae—*Agave horrida* Lem ex Jacobi (photo credit: Etelvina Gándara); Brodiaeoideae—*Bessera elegans* Schult.f. (photo credit: Etelvina Gándara); Nolinoideae— *Beaucarnea gracilis* Lem. (photo credit: Etelvina Gándara); Scilloideae—two unknown *Ledebouria* species (photo credit: Cody Coyotee Howard); Lomandroideae—Lomandra collina (R.Br.) Ewart (photo credit: *Lomandra collina* Goods, Graham & Maree, CC BY-NC-SA 4.0); Asparagoideae—*Asparagus alopecurus* (Oberm.) Malcomber & Sebsebe (photo credit: Philip Bentz).

Asparagaceae contains a diversity of taxa, several of which are horticulturally (e.g., snake plant, *Dracaena* (*Sansevieria*) *trifasciata* (Prain) Mabb.; garden hyacinth, *Hyacinthus orientalis* L.; hosta, *Hosta* Tratt. spp.) and agriculturally important (e.g., garden asparagus, *Asparagus officinalis* L.; tequila and mezcal agaves, *Agave tequilana* F.A.C.Weber, *A. angustifolia* Haw.). Furthermore, there are many iconic, arid-adapted plants including keystone (e.g., Joshua tree, *Yucca brevifolia* Engelm.; Yoder et al. 2024) or endangered species (e.g., Dragon blood tree, *Dracaena draco* (L.) L.; Cartereau et al. 2025) in the group. However, one fairly obscure taxon, *Aphyllanthes monspeliensis* (Aphyllanthoideae), has repeatedly stymied our understanding of the Asparagaceae phylogeny. This rogue taxon has been a major contributor to the often poorly resolved or conflicting topologies across the Asparagaceae phylogenetic backbone; its inclusion in analyses often causes destabilizing effects (Fay et al. 2000; McPherson et al. 2004; Bogler et al. 2006; Seberg et al. 2012; Timilsena et al. 2022) (Fig. 1). Phylogenomic studies excluding *A. monspeliensis* have typically supported an early split in the Asparagaceae, resulting in two major clades composed of three subfamilies each: 1) Asparagoideae + Lomandroideae + Nolinoideae, and 2) Agavoideae + Brodiaeoideae + Scilloideae (Timilsena et al. 2022; Xia et al. 2022; Bentz and Leebens Mack 2024). Relationships among these six larger subfamilies have been relatively congruent across analyses, but relationships among Asparagoideae, Nolinoideae and Lomandroideae remain uncertain (Seberg et al. 2012; Givnish et al. 2018; Timilsena et al. 2022; Bentz and Leebens Mack 2024) (Fig. 1).

All extant Asparagaceae subfamilies originated within the last ∼55 Myr (Ali et al. 2012, 2013; Buerki et al. 2012; Chen et al. 2013; Givnish et al. 2018; Gunn et al. 2020; Meng et al. 2020; Howard et al. 2023; Ji et al. 2023; Bentz et al. 2024) following a series of rapid lineage radiations that occurred early in the evolution of the clade, with subsequent, independent diversifications occurring at different taxonomic scales across the tree (Chen et al. 2013; Jiménez-Barron et al. 2020; Meng et al. 2020; Heyduk et al. 2021; Smith et al. 2021; van Kleinwee et al. 2022; Howard et al. 2023; Bentz et al. 2024). The earliest radiations within the Asparagaceae may have coincided with global cooling since the Eocene-Oligocene transition, which overall resulted in increased global aridity and climatic seasonality (Zachos et al. 2001). One consequence of rapid radiations is the persistence and incomplete sorting of ancestral polymorphisms among diverging populations during speciation events (i.e., ILS), potentially hindering phylogenetic resolution when gene trees disagree with species trees (Maddison and Knowles 2006). Additionally, WGD or polyploidy events, which are well documented in Asparagaceae (McKain et al. 2012; Gunn et al. 2020; Nath et al. 2023), can underlie gene tree-species tree discordance via paralogs inferred as orthologs (e.g., (Xiong et al. 2022). WGDs can provide novel genomic structure for selection to act upon resulting in rapid radiations as well as ecological divergence and phenotypic innovation (McKain et al. 2012; Tank et al. 2015; Stull et al. 2021). Hybridization and introgression, which can promote morphological and ecological diversification (Arnold 1992; Jang et al. 2018; Heyduk et al. 2021), can also drive gene tree-species tree discordance by re-mixing previously assorted loci (Linder and Rieseberg 2004; Crowl et al. 2020). Rapid diversifications, ILS, introgressions, and WGDs are well documented in Asparagaceae and may have uniquely or synergistically spurred the disparate morphological, ecological, and overall species diversification patterns observed across the Asparagaceae phylogeny.

The majority of Asparagaceae taxa inhabit arid or semi-arid, temperate regions across the globe (excluding Antarctica) (Pires and Sytsma 2002; Buerki et al. 2012; Gándara et al. 2014; Norup et al. 2015; Givnish et al. 2018; Heyduk et al. 2023; Ji et al. 2023), but some are also found in more mesic climates (Bogler et al. 2006; Gunn et al. 2020). Overall, no synapomorphy unites the Asparagaceae, but drought adaptive traits (e.g., succulence, underground storage organs, CAM photosynthesis) are fairly common across all seven subfamilies (Givnish et al. 2018). Crassulacean Acid Metabolism or CAM photosynthesis, for example, is well studied within the Agavoideae (Heyduk et al. 2023) and is also documented in the Nolinoideae (Martin et al. 2019; Reyes García et al. 2022). Of particular prevalence across all Asparagaceae subfamilies, however, is belowground bud placement. For example, Scilloideae are all geophytes (i.e., herbaceous, terrestrial plants with belowground buds) with bulbs, Brodiaeoideae are all cormous geophytes, and Asparagoideae and Aphyllanthoideae are mostly rhizomatous (Raunkiaer 1934; Rees 1972). The Agavoideae, Nolinoideae and Lomandroideae contain a relatively larger variety of growth habits across taxa including bulbous geophytes (e.g., *Hesperocallis undulata* A. Gray, Agavoideae), tuberous geophytes (e.g., *Eriospermum* Jacq. ex Willd., Nolinoideae), leafy rhizomatous herbs (e.g., *Hosta*, Agavoideae; *Polygonatum* Mill., Nolinoideae; *Lomandra* Labill., Lomandroideae), above ground succulents (e.g., *Agave* L. spp., Agavoideae; *Dracaena* Vand. ex L. (*Sansevieria* Petagna) spp., Nolinoideae), and arborescent/shrubby plants (e.g., *Yucca brevifolia*., Agavoideae; *Dracaena draco*, Nolinoideae; *Lomandra* spp., Lomandroideae). Root morphology also varies across Asparagaceae with contractile, fibrous and tuberous rooted species found throughout (Gunn et al. 2020; Bentz et al. 2024). The uncertainty of the Asparagaceae phylogenetic backbone, coupled with disparities in sampling and research focus across subfamilies, has limited our ability to comparatively investigate the processes that have contributed to the immense phenomic and genomic diversity found across the Asparagaceae. For example, different sexual systems have evolved in the family, including multiple independent origins of dioecy (i.e., separate male and female plants) across three closely related subfamilies (Asparagoideae, Lomandroideae and Nolinoideae) (Norup et al. 2015; Hernández-Sandoval and Rebman 2018; Bentz et al. 2024; Gunn et al. 2024); however, our understanding of sex chromosome evolution is limited to two species of *Asparagus* (Asparagoideae) (Harkess et al. 2017; Bentz et al. 2026).

Here, we use an Asparagaceae-specific sequence capture probe set targeting 1726 low-copy nuclear loci (Bentz and Leebens Mack 2024), coupled with an extensive generic sampling across the Asparagaceae subfamilies, to resolve evolutionary relationships and examine broadscale, historical biogeographical shifts across the phylogeny. We also leverage discordance patterns across and within various phylogenetic analyses to explore relationships among the seven Asparagaceae subfamilies and unpack evolutionary phenomena underlying gene tree-species tree discordance. Our approach allows us to examine the biological factors underlying discordance across the backbone of Asparagaceae. More broadly, our investigation will lead to a better understanding of the various evolutionary processes that may occur during rapid lineage radiations.

## MATERIALS & METHODS

### Taxon Sampling and Targeted Capture

Since our aims revolved around investigating the drivers of phylogenetic discordance along the Asparagaceae backbone, we aimed for generic-level sampling across Asparagaceae. To do this, we first gathered accepted generic taxonomic concepts of the Asparagaceae from the Plants of the World Online data portal (POWO 2019). We further expanded this list to include genera subsumed into different taxonomic entities (e.g., *Calibanus* Rose [synonymized with *Beaucarnea* Lem.], *Sansevieria* [synonymized with *Dracaena*], *Manfreda* Salisb./*Polianthes* L./*Prochnyanthes* Zucc. [some authors have synonymized these with *Agave*]) (Rojas-Piña et al. 2014; Takawira-Nyenya et al. 2018; Thiede et al. 2019) as well as the resurrected genus *Behria* Greene (Gándara et al. 2024). Within the Scilloideae, we adopted narrower generic concepts, which split large and morphologically complex groups, such as *Ornithogalum* L. and *Drimia* Jacq. ex Wild., into distinct taxonomic entities based on robust molecular phylogenetic and morphological evidence (Martínez-Azorín et al. 2011, 2023; Martínez Azorín et al. 2023). Additionally, we included three Amaryllidaceae taxa and one Asphodelaceae taxon as outgroup representatives (Table S1). Samples (either silica-dried leaf tissue, herbarium voucher tissue, fresh/flash frozen leaf tissue, or DNA extracted material; Table S1) were sent to the University of Minnesota Genomics Center (Minneapolis, MN, USA) for DNA extraction (when applicable), library preparation, and targeted sequencing using an Illumina NovaSeq6000 platform. For targeted sequencing in all samples, we used the Asparagaceae1726 targeted-capture probe set (synthesized by Daicel Arbor Biosciences, Ann Arbor, Michigan, USA), which was developed to improve phylogenetic resolution across the Asparagaceae clade (Bentz and Leebens Mack 2024). The Asparagaceae1726 probe set includes 206 loci from the Angiosperms353 probe set (Johnson et al. 2019) facilitating the integration of our data with other phylogenomic studies. Lastly, we included 24 already available samples from Bentz and Leebens Mack (2024), and eight shotgun sequence samples for Agavoideae species (Table S1) to further expand taxon representation.

### Sequence Cleaning, Assembly and Alignment

Sequencing reads were cleaned with fastp v.0.23.2 (Chen et al. 2018), then mapped to the reference targets (i.e., nucleotide coding sequences) and assembled using HybPiper v.2.3.1 with the BWA option for read mapping (Johnson et al. 2016). For read mapping, we used the Asparagaceae1726 nucleotide reference fasta file v.2.0 (https://github.com/bentzpc/Asparagaceae1726), which reduces paralogs and increases on-target ortholog recovery in silico, compared to previous versions. Multiple sequence alignments for each target locus were estimated using MAFFT v.7.520 and the -auto flag (Katoh and Standley 2013). Poorly aligned sequences were trimmed using trimAl v.1.4.1 using the *-automated1* flag (Capella-Gutiérrez et al. 2009).

Additionally, we extracted off-target chloroplast reads from all samples to assemble 77 chloroplast protein coding genes for phylogenetic analysis. HybPiper v.2.3.1 (Johnson et al. 2016) was used, with the same methods as above, to map and assemble the chloroplast genes. However, instead of using supercontigs (i.e., exon+intron assemblies) we used only the exon assemblies for further analysis in order to avoid long introns and highly gapped alignments. Reference chloroplast genes used for read mapping were from *Asparagus schoberioides* Kunth (NCBI GenBank: NC_035969) and *A. aethiopicus* L. (NCBI GenBank: MZ337394). All resulting chloroplast gene assemblies were separately aligned with MAFFT v.7.520 and trimmed with trimAl v.1.4.1, as before, except using more stringent parameters with MAFFT (--localpair, --maxiterate 1000). The clean assemblies were then concatenated into a supermatrix and analyzed as a single locus.

### Phylogenetic Reconstruction

We took multiple approaches to more fully investigate phylogenetic discordance along the Asparagaceae backbone. Using the Asparagaceae1726 dataset, phylogenetic reconstruction was first performed using maximum likelihood (ML) as implemented in IQ-Tree v.2.1.2 (Minh et al. 2020). We performed a partitioned analysis with a concatenated supermatrix using the *-p* command, the GENE partition resampling scheme (--sampling GENE) (Nei et al. 2001), with automatic selection of the best fit substitution model and partitioning scheme followed by tree reconstruction (-m TESTMERGE) (Lanfear et al. 2012). To assess branch support, we performed both 1000 ultrafast bootstrap (UFBoot) approximations (-B 1000) (Hoang et al. 2018) as well as the SH-like approximate likelihood ratio test (SH-aLRT) (Guindon et al. 2010) with 1000 bootstrap replicates (-alrt 1000). We considered a branch strongly supported if UFBoot and SH-aLRT were >=95% and >=80%, respectively.

Using IQ-Tree v.1.6.12 (Nguyen et al. 2015) with automatic model selection (-m MFP) (Kalyaanamoorthy et al. 2017), we reconstructed two separate ML trees using the chloroplast dataset: one using 1000 UFBoots, and another using 100 standard (nonparametric) bootstraps (BS). In single gene trees (i.e., both plastome trees in this case), UFboot support is less biased and should roughly correspond to posterior probabilities, thus reliable branches should exhibit support >=95% (Minh et al. 2013).

For estimating species relationships, a coalescent-based, gene tree/quartet summary method, rather than a concatenated supermatrix of unlinked loci is generally preferred because concatenation-based approaches implicitly assume an absence of recombination and ILS (i.e., that a single history is shared amongst all genes). This can result in statistical inconsistencies in multi-locus datasets with high levels of gene tree discordance (Kubatko and Degnan 2007; Edwards et al. 2016). Coalescent-based summary methods (e.g., ASTRAL), however, assume free recombination between genes and account for gene tree discordance due to ILS (Zhang et al. 2017). IQ-Tree v.1.6.12 was used for ML gene tree inference with 1000 UFBoots and automatic model selection (-m MFP). To estimate the species tree, we used weighted ASTRAL (wASTRAL) v.1.22.3.7 (Zhang and Mirarab 2022), which is a reimplementation of ASTRAL-III (Zhang et al. 2017) that accounts for phylogenetic uncertainty by integrating signals from branch length and branch support in gene trees. Gene tree branches with <10% UFBoots support were collapsed into polytomies for a separate ASTRAL-III analysis, replicating previous species tree methods for comparison with our expanded taxon dataset (Bentz and Leebens Mack 2024). Clade support was assessed according to local posterior probabilities (LPP) and other quartet-based metrics, as outlined below. All resulting trees were rooted with *Dianella tasmanica* Hook.f. (Asphodelaceae).

### Phylogenetic Discordance

Using the wASTRAL tree as the main topology, we assessed gene tree quartet concordance vs. discordance among all nuclear gene trees as implemented with Quartet Sampling (QS) v.1.3.1 (Pease et al. 2018) using RAxML-ng v.1.2.2 (Kozlov et al. 2019) as the likelihood evaluation engine. QS was performed with conservative parameters including 1000 replicates and a likelihood threshold of 2.0; Quartet Concordance (QC), Quartet Differential (QD) and Quartet Informativeness (QI) were used to interpret gene tree discordance (Pease et al. 2018). Additionally, discordance was assessed based on quartet frequency support for the main, most likely topology (*q1*) and two alternative topologies (*q2* and *q3*) reported by ASTRAL (Zhang et al. 2017; Zhang and Mirarab 2022). We then also used DiscoVista (Sayyari et al. 2018) to further quantify and visualize discordance across the nuclear gene trees using the relative frequency of gene tree quartets.

We used Quaint v.1.0.0 (Baldwin and Leebens-Mack 2026) (https://github.com/ethan-baldwin/quaint) to test whether introgression may be implicated in the observed gene tree discordance across the Asparagaceae phylogeny. This method employs quartet-based statistical tests similar to the ABBA-BABA test to identify signatures of introgression. The analysis quantifies skew in alternate gene tree topologies by calculating a statistic (Dq) similar to Patterson’s D (Green et al. 2010) for a quartet of taxa and assesses significant skew with a chi-square test. These statistics are summarized for each pair of taxa by taking the mean Dq of all quartets while setting non-significant quartets to 0. Inputs for Quaint included the collection of gene trees used for wASTRAL and a fixed species tree topology (here being the wASTRAL topology). We used *Dianella tasmanica* to root gene tree quartets. Results from Quaint were summarized with mean Dq plotted in a matrix—any shade of red indicates statistical significance of ABBA>BABA.

### Divergence Time Estimation

We performed divergence time estimation using penalized likelihood as implemented in treePL (Smith and O’Meara 2012) for each topology recovered (i.e., ML nuclear, ML cpDNA, wASTRAL). However, rather than estimating divergence times on one phylogeny only, we took an iterative approach to generating uncertainty around node age estimates (see https://github.com/sunray1/treepl). First, we generated 100 bootstrap alignments from the nuclear locus alignment using RAxML v.8.2.12 (Stamatakis 2014), incorporating the best-fitting partitioning scheme output from the respective IQ-Tree analysis (note: an independent ML analysis was performed using the wASTRAL topology as a constraint to obtain the best partitioning scheme). We then performed tree reconstruction using a GTRGAMMA nucleotide substitution model and constrained each analysis to either the ML topologies (ML nuclear, ML cpDNA) or the wASTRAL topology. This resulted in 100 phylogenies for each topology, which were then each input into treePL for divergence time estimation. For each phylogeny, we performed a priming step followed by cross validation with the settings cvstart=100000, cvstop=0.000001, and cvmultstep=0.09, and this was repeated three separate times for each individual tree. From the cross validation results, each phylogeny had the best smoothing value applied as determined by a chi-square analysis. During the whole process, we applied two ingroup fossil calibrations (Iles et al. 2015): 1) the Agavoideae crown node (minimum 14.5 Ma), and 2) the Lomandroideae crown node (minimum 22 Ma). We also placed one outgroup fossil calibration at the Amaryllidaceae crown node (minimum 48.88 Ma) (Pigg et al. 2018). We also placed two secondary calibrations (Givnish et al. 2018): 1) the split between Asphodelaceae and Asparagaceae (52.3–58.1 Ma), and 2) the Asparagaceae crown node (47.0–51.9 Ma). After each individual topology was calibrated, we summarized the resulting 100 ultrametric trees using TreeAnnotator v.2.7.7 (Bouckaert et al. 2014) to calculate median age estimates and 95% confidence intervals.

### Biogeographical Analysis

To investigate the timing and geographic regions of historical overlap among the major Asparagaceae clades, we used BioGeoBEARS v.1.1.3 (Matzke 2013a) to reconstruct the estimated ancestral range at each node of the phylogeny. We categorized the distribution of each tip in the phylogeny at the generic level using a semi-continental (i.e., “*botanical continents*”) coding scheme using the World Geographical Scheme for Recording Plant Distributions (Brummitt et al. 2001). When coding tips, we collapsed monophyletic genera and/or synonyms that shared the same biogeographical distribution. We tested three biogeographical models (DEC, DIVA-LIKE and BAYAREA-LIKE) both with and without the jump dispersal (+j) parameter included (Matzke 2013b, 2014). The best biogeographic model was determined using Akaike information criterion (AIC).

## RESULTS

### Sequences

In total, 151 samples were included in our analyses. We aimed to sample 143 Asparagaceae genera, and successfully obtained 114 genera (i.e., 80% generic coverage) and 144 different Asparagaceae species (Table S1). All but one of the 1726 possible target loci were recovered in >1 sample. Seventy percent (1215/1726) of the target loci were recovered in at least 135 samples, or 89%, of the 151 total samples, and 85% of the total target loci were recovered in at least 100 samples. One target locus (target 1702) was recovered in only one sample and was excluded from downstream analysis, while all other targets were utilized downstream, even those with relatively poor recovery rates across samples (i.e., 2% of the targets were recovered in <20 samples). Since ASTRAL breaks gene trees into four-leaf quartets, we deemed all gene trees with >4 tips as informative and thus, all target loci that were recovered in >4 samples were included in the species tree analyses.

Of the 151 individuals included, 106 (plus three outgroup taxa) were newly sequenced for the first time using the Asparagaceae1726 targeted capture approach. Of the newly sequenced samples (which did not include Asparagoideae), the average number of mapped reads ranged from 3,563,332 (Scilloideae) to 10,159,682 (Aphyllanthoideae), the average percent on target reads ranged from 23.4% (Aphyllanthoideae) to 49.2% (Lomandroideae), the average total bases recovered ranged from 1,283,167 (Scilloideae) to 1,680,675 (Aphyllanthoideae), and the number of paralog warnings ranged from 8.8 (Lomandroideae) to 77.5 (Aphyllanthoideae) (Fig. S1; Table S2). Note that the monotypic Aphyllanthoideae is represented by six samples, which overinflates sequencing statistics. For the chloroplast data, at least 43 chloroplast genes were recovered from each sample and >95% of the total samples yielded >70 chloroplast genes (out of the 77 selected chloroplast genes) (Table S2).

### Relationships among Agavoideae, Aphyllanthoideae, Brodiaeoideae and Scilloideae

All six samples of *Aphyllanthes monspeliensis* (Aphyllanthoideae) formed a strongly supported clade in all analyses (LPP=1.0, UFBoot=100) (Fig. S2, S3, S4). In the wASTRAL analysis, Aphyllanthoideae was strongly supported in a clade with Agavoideae + Brodiaeoideae + Scilloideae (LPP=1) with 47% of gene tree quartets supporting this topology (*q1*=47%), compared to the top two alternatives (*q2*=27%, *q3*=26%), and resulted in slightly more discordant than concordant quartets (QC=-0.01, QD=0.83, QI=0.26) (Fig. S5). DiscoVista reported a majority of gene tree quartets supporting a monophyletic Agavoideae + Brodiaeoideae + Scilloideae (t1, 1565 gene trees out of 1619 gene trees) (Fig. 2D, Panel 8), with an almost equal frequency between topologies t2 and t3 (435 vs 448 gene trees, respectively). A majority of gene tree quartets placed Aphyllanthoideae in a clade with Agavoideae + Brodiaeoideae + Scilloideae (t1, 643 gene trees out of 1536 gene trees), whereas the next highest of 496 gene tree quartets (t3) placed Aphyllanthoideae with the outgroups (i.e., as sister to all remaining Asparagaceae) (Fig. 2D, Panel 9). Branch support for an Aphyllanthoideae + Agavoideae + Brodiaeoideae + Scilloideae clade were similar in the unweighted ASTRAL-III analysis, though with skewed support for *q2* > *q3* (*q1*=42%, *q2*=32%, *q3*=26%) (Fig. S6). The Agavoideae + Brodiaeoideae + Scilloideae clade was also strongly supported (LPP=1) and showed little discordance among quartets (wASTRAL: *q1*=95%, *q2*=3%, *q3*=2%; QC=0.042, QD=0.98, QI=0.26) (ASTRAL-III: *q1*=44%, *q2*=29%, *q3*=28%) (Fig. S2). According to an ASTRAL-III polytomy test, polytomies were rejected (*p*=0) across the entire Asparagaceae backbone including for both of the aforementioned nodes. According to the concatenation ML analysis of nuclear genes, the six larger subfamilies, not including Aphyllanthoideae, formed well-supported clades (UFBoot=100, SH-aLRT=100), but the placement of Aphyllanthoideae as sister to all remaining taxa received low support (UFBoot=54.3, SH-aLRT=60) (Fig. 2). In the chloroplast phylogeny, Aphyllanthoideae was placed sister to Brodiaeoideae + Scilloideae with low support (BS=76, UFBoot/SH-aLRT=71.2/86), but nested within a strongly supported clade of Agavoideae + Brodiaeoideae + Scilloideae (BS=100, UFBoot/SH-aLRT=100/100) (Fig. 2). Lastly, Quaint revealed little evidence of introgression between Aphyllanthoideae and other subfamilies (mean Dq ∼0) (Fig. 3, S7) suggesting that the observed gene tree-species tree discordance was likely caused by rapid diversification driving ILS. Quartet concordance for the placement of Aphyllanthoideae was lower in the concatenation-based species tree (QC=-0.064 for a sister relationship between Aphyllanthoideae and the rest of Asparagaceae, Fig. S8) compared to the wASTRAL tree (QC=-0.01 for Aphyllanthoideae forming a clade with Agavoideae + Brodiaeoideae + Scilloideae, Fig. S5). Various species relationships within major clades showed low quartet concordance values, which may call for population-level investigation to elucidate further (Fig. S5). Nonetheless, relationships among Agavoideae, Brodiaeoideae, and Scilloideae were concordant across all analyses in this study (Fig. 2) and agree with previous studies (Fig. 1).

**Figure 2.**
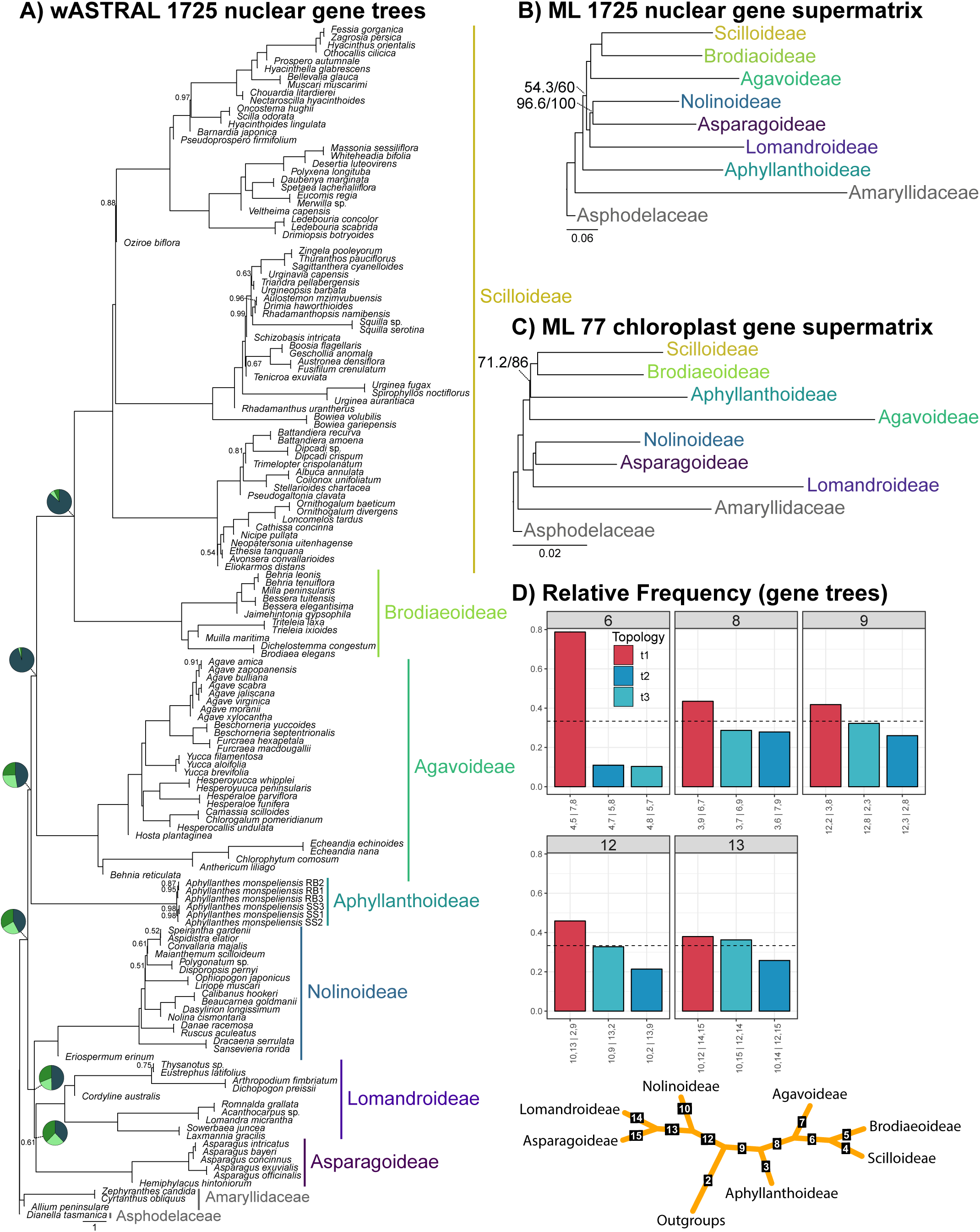
Phylogenomic relationships among the seven Asparagaceae subfamilies across datasets and analyses. A) Species relationships using multispecies coalescent methods (wASTRAL) and 1775 nuclear locus trees. Support values show only those nodes below a local posterior probability (LPP) equal to 1.0. Pie charts corresponding to major backbone branches represent alternative quartet frequencies from wASTRAL: dark green represents the percent of quartets supporting the main topology, or *q1* (as illustrated here), while medium green (*q2*) and light green (*q3*) represent the top two alternative topologies. B) Species relationships analyzed using maximum likelihood and a concatenated, partitioned matrix of 1775 nuclear loci. Support values shown for branches with less than 100% UFBoot or SH-aLRT support (UFBoot / SH-aLRT). C) Phylogenetic relationships analyzed using maximum likelihood and a concatenated matrix of 77 chloroplast genes. Support values shown as described above (UFBoot / SH-aLRT). D) Quartet frequencies supporting major clades across the Asparagaceae phylogenetic backbone estimated with DiscoVista. The main topology (t1) is shown in red, and the alternative two topologies (t2 and t3) are shown in shades of blue. The dotted line marks a ⅓ threshold of the total quartets supporting a single topology.

**Figure 3.**
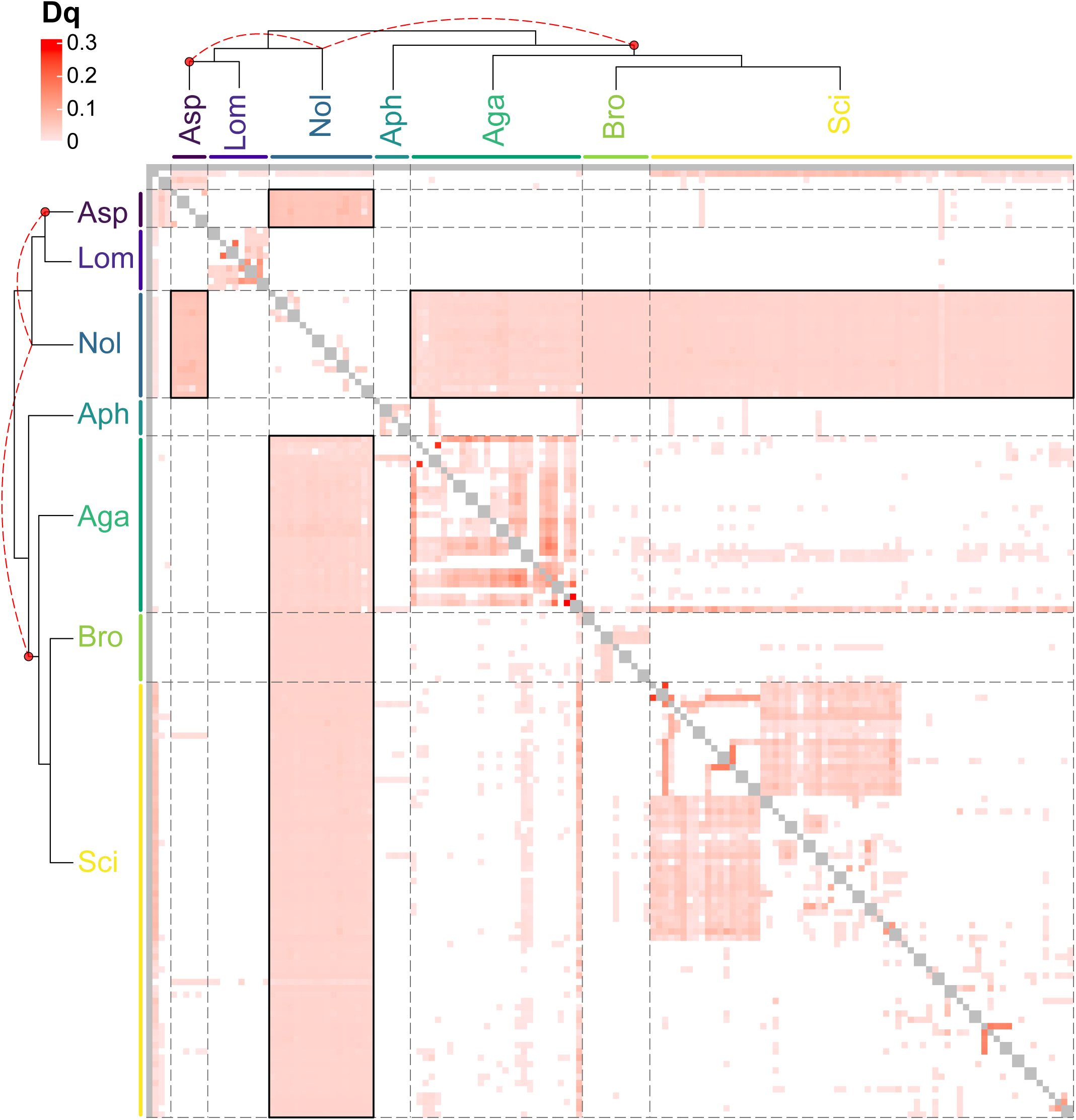
Heatmap summarizing a quartet-based ABBA-BABA analysis performed with Quaint to test for historical introgressions across the Asparagaceae phylogeny. Each tile displays the mean Dq, which quantifies skew in alternate quartet topologies similar to Patterson’s D, for a pair of taxa. Any value of Dq greater than zero may represent evidence of introgression between two branches. The bold rectangles highlight significantly elevated Dq between entire subfamilies/clades; indicating that a possible ancient hybrid event, involving an ancestor of the Asparagoideae (Asp) clade and the Agavoideae (Aga) + Brodiaeoideae (Bro) + Scilloideae (Sci) clade, may have contributed to the origin of the Nolinoideae clade. Abbreviations for remaining subfamilies are Aphyllanthoideae (Aph) and Lomandroideae (Lom). Tree topology is from Weighted ASTRAL. Red dotted lines show the hypothesized ancient reticulation between ancestral backbone branches implicated in the origin of Nolinoideae. Phylogeny with tip labels included can be found in Figure S7.

### Relationships among Asparagoideae, Lomandroideae and Nolinoideae

The Asparagoideae, Lomandroideae and Nolinoideae formed a well-supported clade in all analyses (LPP=1.0, UFBoot/SH-aLRT=100/100) with ∼50% (*q1*) of the wASTRAL quartet gene trees supporting the species tree topology (Fig. S2, S3, S4). However, we found discordance across analyses regarding inter-subfamilial relationships. In both the wASTRAL and ASTRAL-III analyses, Nolinoideae was recovered as sister to an Asparagoideae + Lomandroideae clade with 0.61 and 0.98 LPP support, respectively (Fig. 2, S6). Interestingly, skewed support for alternative quartet topologies were observed with wASTRAL for the Asparagoideae + Lomandroideae + Nolinoideae clade (*q2*=19%, *q3*=31%, QC=0.06, QD=0.6, QI=0.23) as well as the Asparagoideae + Lomandroideae sister relationship (*q1*=38%*, q2*=24%, *q3*=38%, QC=0.016, QD=0.8, QI=0.25) recovered in the wASTRAL tree (Fig. S5); similarly skewed quartet frequencies were also found in the ASTRAL-III analysis (Fig. S6). In the ML concatenated species tree and ML chloroplast tree, Lomandroideae was recovered as sister to an Asparagoideae + Nolinoideae clade with decent support in the former (UFBoot/SH-aLRT=96.6/100) and strong support in the latter (BS/UFBoot/SH-aLRT=100) (Fig. 2). Regarding the placement of Nolinoideae, quartet concordance was lower in the concatenation-based species tree (QC=-0.031 for a Nolinoideae + Asparagoideae clade, Fig. S8) compared to the wASTRAL tree (QC=0.016 for a Lomandroideae + Asparagoideae clade, Fig. S5). The DiscoVista gene tree quartet frequency analysis showed that out of 1608 gene trees, 737 (t1) placed Nolinoideae with Asparagaceae + Lomandroideae, whereas 526 gene trees placed Nolinoideae with Aphyllanthoideae + Agavoideae + Brodiaeodieae + Scilloideae (t3) (Fig. 2D, Panel 12). We also found an almost equal frequency between topologies t1 and t3 (632 vs 604 gene trees out of 1666 gene trees, respectively), but not t2 (429 gene trees), for the Asparagoideae + Nolinoideae + Lomandroideae relationships (Fig. 2D, Panel 13). Interestingly, in our quartet-based ABBA-BABA test, we found significant signatures of introgression between all members of Nolinoideae and Asparagoideae (mean Dq∼0.05) as well as between Nolinoideae and almost all members of the larger Agavoideae + Brodiaeoideae + Scilloideae clade (mean Dq∼0.03) (Fig. 3, S7). While Quaint tests for introgression between two tips of a species tree, similar mean Dq values between all taxa within two clades is expected when ancient introgression between the ancestors of the respective clades occurs.

### Divergence times

Median divergence time estimates placed the origin of the Asparagaceae sometime within the past ∼50 Myr. Across the three topologies (ML nuclear, ML cpDNA, wASTRAL), divergence time estimates largely overlapped with one another (Table 1, Table S3). The wASTRAL topology produced generally younger median estimates for many of the subfamily crown nodes (Table 1, Table S3). The ML nuclear topology estimated that Lomandroideae diverged from Asparagoideae + Nolinoideae ∼46.6 Ma (95% CI 46.8–40.9), whereas in the wASTRAL topology Nolinoideae diverged from Asparagoideae + Lomandroideae ∼45.2 Ma (95% CI 50.67–45.0).

**Table 1.** Divergence times and reconstructed biogeographic ranges for each Asparagaceae subfamily. Stem and crown ranges reflect the 95% confidence intervals from each separate treePL analysis for each topology: maximum likelihood—nuclear loci (ML nuclear), maximum likelihood—chloroplast loci (ML cpDNA), and Weighted ASTRAL (wASTRAL).

Age estimation across topologies for the Aphyllanthoideae crown and stem were similar (Table S3). The ML nuclear topology, which reconstructed Aphyllanthoideae as sister to all remaining Asparagaceae taxa, estimated a median stem age of 48.9 Myr (95% CI 49.0–47.0) and a median crown age of 2.4 Myr (95% CI 2.4–1.9). The wASTRAL topology, which placed Aphyllanthoideae sister to a clade of Agavoideae + Brodiaeoideae + Scilloideae, estimated a median stem age of 46.32 Myr (95% CI 51.3–46.1) and a median crown age of 2.7 Myr (95% CI 2.9–2.4). Lastly, the ML chloroplast topology, with Aphyllanthoideae sister to Brodiaeoideae + Scilloideae, estimated the Aphyllanthoideae stem at 48.7 Myr (95% CI 50.8–46.0) and crown at 0.30 Myr (95% CI 0.8–0.2), the latter being the youngest crown age estimates of the three topologies.

### Asparagaceae biogeographic history

For each of the three topologies, the best biogeographic model as determined by AIC was BAYAREA-LIKE, with further model improvement when including the +j parameter (Table S4). For each topology, Africa was the reconstructed historical range for the Asparagaceae (Table 1, S3; Figs. S9, S10, S11). Brodiaeoideae and Lomandroideae were favored to have originated in North America and Australia, respectively, whereas Africa was favored for the remaining subfamilies (Table 1). Regardless of phylogenetic placement, Aphyllanthoideae was favored to have originated in Africa (Table 1). Minor differences in the probability of occurrence for the Asparagaceae backbone were found across topologies, yet the estimated ancestral range was consistent across topologies for each subfamily (Table 1, S3).

## DISCUSSION

In this study we used the most comprehensive sampling of Asparagaceae genera to date, coupled with chloroplast sequences and targeted sequencing of 1726 low-copy nuclear genes, to resolve evolutionary relationships and investigate sources of gene tree discordance across the phylogenetic backbone of the family. To further explore gene tree-species tree discordance, we compare results from concatenation-based vs. coalescent-based species tree methods, along with a separate chloroplast phylogeny. Concatenation-based phylogenetic analysis (i.e., using a supermatrix of gene sequences) inherently assumes that each concatenated gene in a sequence matrix shares the same evolutionary history, which can often lead to spurious topology resolutions in the presence of rapid diversifications and ILS (Warnow 2015; Mendes and Hahn 2018). However, a multispecies coalescent approach may yield more accurate species relationships when ILS is present, by leveraging frequency measurements of discordant gene tree topologies. Here, we performed multispecies coalescent summary methods using Weighted ASTRAL (wASTRAL), which downweights long terminal branches and poorly supported nodes, further reducing the impact of long branch attraction and uncertainty surrounding short branches in individual gene trees (Zhang and Mirarab 2022). Importantly, wASTRAL outperforms the unweighted ASTRAL algorithm and is more accurate than concatenation-based approaches in rapidly diversifying clades with short internodes (Zhang and Mirarab 2022).

At the broadest scale, the multispecies coalescent tree (Fig. 2A) and chloroplast phylogeny (Fig. 2C) both show two major clades dating back to the earliest lineage diversifications in the family (∼45–40 Ma, Tables 1, S3): 1) Asparagoideae + Lomandroideae + Nolinoideae; and 2) Aphyllanthoideae + Agavoideae + Brodiaeoideae + Scilloideae. Across the backbone of the Asparagaceae phylogeny, gene tree discordance patterns suggests that ILS— resulting from rapid lineage diversifications in Africa ∼49 Ma—explains much of the observed conflicting topologies, including the placement of Aphyllanthoideae (Fig. 2). Our comprehensive generic-level sampling coupled with dense molecular sampling enabled robust inference for the placement of Aphyllanthoideae within the Asparagaceae family. Aphyllanthoideae is strongly supported as sister to a larger clade with Agavoideae, Scilloideae and Brodiaeoideae (Fig. 2A); however, gene tree discordance frequencies suggest that an ancient reticulation involving an Aphyllanthoideae ancestor may further explain its instability in phylogenetic analysis, beyond ILS. Additionally, quartet-based ABBA-BABA tests (Fig. 3) revealed that ancient introgression between two divergent ancestral lineages may underlie the discordant resolutions for Nolinoideae within the Asparagoideae + Lomandroideae + Nolinoideae clade.

### Incomplete lineage sorting, and the origin of Aphyllanthoideae

The inclusion of *Aphyllanthes monspeliensis* (Aphyllanthoideae) in phylogenetic analysis has historically led to poor resolution among the Asparagaceae subfamilies (Fig. 1), which was also observed in our analysis of concatenated nuclear genes (Fig. 2B) and chloroplast tree (Fig. 2C). This discordance has been hypothesized to be driven by technical (e.g., long-branch attraction), biological (e.g., historical introgression, ILS) and/or lineage-specific (e.g., WGDs) factors. Aphyllanthoideae arose following rapid diversification(s) of ancestral Asparagaceae lineages, as marked by the extremely short branches along the backbone of the phylogeny (Fig. 2A). Earlier phylogenomic work of 602 nuclear genes resolved Aphyllanthoideae as the sister lineage to Agavoideae + Brodiaeoideae + Scilloideae, but with very poor support (LPP = 0.22) (Timilsena et al. 2022). In our wASTRAL analysis, we found strong support for this relationship (LPP=1.0, Fig. 2A), as indicated by 1) no skew in alternative quartet topology frequencies supporting the placement of Aphyllanthoideae using local branch tests (Fig. 2A, pies along backbone) and 2) the majority of gene tree (clade-based) quartet frequencies supporting the top two alternative topologies each being less than one-third (Fig. 2D). ABBA-BABA tests showed no evidence of historical introgressions involving Aphyllanthoideae and an ancestral backbone branch (i.e., the Most Recent Common Ancestor (MRCA) of a subfamily) within the Asparagaceae (Fig. 3). However, using DiscoVista to measure gene tree topology frequencies among major clades, we found considerable skew favoring one alternative topology suggesting possible reticulations between an Aphyllanthoideae ancestor and an extinct Asparagaceae backbone branch, or a lineage preceding the origin of the family (i.e., Fig 2D, panel 9). Additional taxon sampling from Amaryllidaceae and/or Asphodelaceae is required to test the possibility of more ancient reticulations. Regardless, our plastid phylogeny (Fig. 2C) agrees with the coalescent-based species trees (Fig. 2A, S6) in they both firmly place Aphyllanthoideae in a clade with Agavoideae, Brodiaeoideae and Scilloideae. An additional source of gene tree discordance may result from the inclusion of paralogs in phylogenetic analysis. However, the Asparagaceae1726 probe set was specifically designed to select against the inclusion of paralogs, and the limited number of paralogs that may result in Asparagaceae1726 datasets do not lead to topology differences (Bentz and Leebens Mack 2024). Recent work has also shown that the inclusion of paralogs may not be as detrimental in phylogenomic analysis of hundreds to thousands of genes and sometimes can even improve resolution (Ufimov et al. 2022; Yan et al. 2022).

Aphyllanthoideae diverged from its MRCA ∼46.3 Ma in Africa, soon after the origin of the family (Table 1, S3). Today, the monotypic *Aphyllanthes monspeliensis* is geographically restricted to the western Mediterranean Basin, overlapping in distribution with many other Asparagaceae taxa (e.g., *Asparagus*, many Scilloideae and Agavoideae species). The ecological and geological processes leading to the now geographically restricted nature of *A. monspeliensis* are unknown, but our sampling of multiple individuals from two distinct populations suggests a fairly young origin for the species (< 3 Ma) (Table S3). Interestingly, the number of paralog warnings (i.e., multiple, long contigs per gene found) were elevated in our *A. monspeliensis* samples (Fig. S1, Table S2), suggesting historical, lineage-specific duplication events may have occurred in Aphyllanthoideae. Perhaps Quaternary glaciation cycles in Europe, along with the onset of major climatic oscillations in southern Europe and northern Africa during the Pliocene-Pleistocene (Rundel et al. 2016), synergistically played a role in constraining dispersal of *A. monspeliensis*, as well as driving other possible *Aphyllanthes* lineages to extinction. Future population genomics work should explore population structure within *A. monspeliensis* to further understand the extant diversity present in the species.

### Ancient hybridization and the origin of Nolinoideae

The Nolinoideae clade is commonly resolved as sister to Asparagoideae in analyses of few molecular markers (Fig. 1). However, in phylogenomic analyses, Nolinoideae has alternatively been recovered as sister to an Asparagoideae + Lomandroideae clade according to both concatenation-(602 nuclear loci; Timilsena et al. 2022) and coalescent-based inference (1,726 nuclear loci; Bentz and Leebens Mack 2024). Analyses performed in our study were similarly discordant with conflicting support for an Asparagoideae + Lomandroideae clade in coalescent-based trees (Fig. 2A, S6), versus an Asparagoideae + Nolinoideae clade in trees inferred from concatenated nuclear genes (Fig. 2B) and chloroplast genomes (Fig. 2C). Whereas most individual gene tree quartets supported an Asparagoideae + Lomandroideae sister relationship (Fig. 2A, D), skewed support for alternative quartet topologies suggests that confounding biology (other than ILS) may be a greater source of discordance among these three subfamilies (Pease et al. 2016; Sayyari et al. 2018). Our quartet-based ABBA-BABA test suggest that an ancient hybridization event, involving the MRCA, or other ancestral lineages, of Asparagoideae and an ancestor of the Agavoideae + Scilloideae + Brodiaeoideae clade, may have contributed to the origin of the Nolinoideae subfamily (Fig. 3). Given the overlapping divergence times and geographic origins of most Asparagaceae subfamilies in Africa (Fig. S9, S10, S11; Table 1, S3), instances of reticulation among the young, ancestral lineages is possible considering the prevalence of deep reticulations across the angiosperm tree of life (reviewed in Stull et al. (2023)). An ancient hybridization hypothesis for the origin of Nolinoideae—wherein introgression between ancestral lineages of Asparagoideae and the Agavoideae + Scilloideae + Brodiaoideae clade led to formation of a budding lineage—could explain the low topological support, gene tree-species tree discordance, and disagreement between nuclear gene concatenation-(Fig. 2B) and coalescent-based (Fig. 2A) reconstructions. Interestingly, chloroplast tree topologies always show strong support for an Asparagoideae + Nolindoideae sister relationship (Fig. 1, 2C) (Bentz and Leebens-Mack 2024) suggesting the ancestral Asparagoideae chloroplast was captured during the putative hybridization event. We cannot outright reject the possibility that an ancestrally shared WGD, followed by lineage-specific genome evolution (e.g., biased gene retention, subfunctionalization, etc.), does not underlie any of the observed gene tree discordance. However, an ancestrally shared WGD should not impact our hybrid origin hypothesis for Nolinoideae because we would expect similar signals of introgression or discordance to be recovered in other subfamilies like Lomandroideae or Aphyllanthoideae.

## CONCLUSIONS AND FUTURE DIRECTIONS

This study opens doors to new research avenues for this economically and ecologically important plant family. The wealth of data provided by this study should be harnessed to further expand understanding of Asparagaceae diversity, from molecular processes to morphological evolution and ecology. Our findings highlight intriguing patterns worthy of additional investigation, such as the underlying mechanisms driving the evolution of dioecy, which has evolved repeatedly in only one of the two major Asparagaceae subclades. Additionally, the diversity of photosynthetic pathways should be assessed across Asparagaceae subfamilies. CAM photosynthesis has evolved repeatedly within Agavoideae and Nolinoideae, both of which are implicated in our Nolinoideae hybridization hypothesis. Could historical hybridization be the cause for these two distantly related lineages sharing this photosynthetic pathway or is CAM photosynthesis more prevalent across the family? Future work incorporating genomic, transcriptomic, and physiological assays across Asparagaceae species should be carried out to fully investigate the evolution of these two important ecological and agricultural traits.

## AUTHOR CONTRIBUTIONS

**Philip C. Bentz** - Conceptualization, Data Curation, Formal Analysis, Acquisition, Resources, Visualization, Writing - original draft, Writing - review & editing; **Ethan Baldwin** - Formal Analysis, Resources, Writing - review & editing; **Mario Martínez-Azorín** - Resources, Writing - review & editing; **Kálmán Könyves** - Resources, Writing - review & editing; **Timothy S. Harvey** - Resources, Writing - review & editing; **Joanne Birch** - Resources, Writing - review & editing; **Etelvina Gándara** - Resources, Writing - review & editing; **Bryan MacNeill** - Resources, Writing - review & editing; **Andrew Salywon** - Resources, Writing - review & editing; **Xiaohan Yang** - Resources, Writing - review & editing; **James Reed** - Resources, Writing - review & editing; **Sierra Jaeger** - Resources, Writing - review & editing; **James H. Leebens-Mack** - Conceptualization, Resources, Writing - review & editing; **Cody Coyotee Howard** - Conceptualization, Data Curation, Formal Analysis, Funding Acquisition, Resources, Visualization, Writing - original draft, Writing - review & editing

## ACKNOWLEDGEMENTS

Samples were contributed from the collections of: The University of Georgia Herbarium (GA), Herbario del Instituto de Botánica (IBUG), Universidad de Guadalajara, Universidad de Alicante (ABH herbarium and Scilloideae living collection), Royal Horticultural Society Garden Wisley, and the private collection of John David. Computing resources were provided by the Oklahoma State University High Performance Computing Center, the University of Central Florida Advanced Research and Computing Center, and Georgia Advanced Computing Resource Center at University of Georgia. The Department of Environment and Nature Conservation of Northern Cape Province (collecting and export permits FLORA046/2010, FLORA047/2010, FLORA069/2011, FLORA070/2011, FLORA61/2/2015, FLORA61/2/2015, FLORA0057/2017, FLORA0058/2017, FLORA0059/2022, FLORA0060/2022), CapeNature of Western Cape Province of South Africa (collecting and export permits AAA005-00080-0028, AAA008-00031-0028, 0027-AAA008-00699, 0028-AAA008-00203, CN35 28 21389, CN17 28 21605), Ezemvelo-KZN Wildlife (collecting permit OP 3969/2015) provided permission to collect specimens. The Ministry of Environment and Tourism of Namibia provided a research/collecting permit 2192/2016. This work was partly supported by H2020 Research and Innovation Staff Exchange Programme of the European Commission, Project 645636: ‘Insect-plant relationships: insights into biodiversity and new applications’ (FlyHigh) and the complementary supporting funds UAUSTI17-03, UAUSTI19-08, ACIE17-01, ACIE18-03 and ACIE21-01 (University of Alicante, Spain).

## DATA AVAILABILITY

Sequencing reads will be made available prior to publication on NCBI’s Sequence Read Archive (SRA submission: SUB16489649) BioProject TBD. The Asparagaceae1726 target files are on GitHub https://github.com/bentzpc/Asparagaceae1726. Data analysis and result files can be found in Dryad (DOI).

## SUPPLEMENTARY MATERIALS

**Figure S1**. Locus summary statistics for the Asparagaceae1726 nuclear loci dataset.

**Figure S2**. Weight ASTRAL topology using gene trees (wASTRAL). Support values are local posterior probability.

**Figure S3**. IQ-Tree topology using concatenation of nuclear loci (ML nuclear). Support values are ultrafast bootstrap support and SH-aLRT (UFBoot / SH-aLRT).

**Figure S4**. IQ-Tree topology using chloroplast data analyzed as one locus (ML cpDNA).

**Figure S5**. Quartet Sampling using the Weight ASTRAL (wASTRAL) topology as input.

**Figure S6**. Phylogenetic reconstruction using ASTRAL-III.

**Figure S7**. Full Quaint figure with tip labels included.

**Figure S8**. Quartet Sampling using the maximum likelihood nuclear (ML nuclear) topology as input.

**Figure S9**. BioGeoBEARS BAYAREALIKE+J on Asparagaceae Weighted ASTRAL (wASTRAL) topology.

**Figure S10**. BioGeoBEARS BAYAREALIKE+J on Asparagaceae maximum likelihood nuclear gene supermatrix (ML nuclear) topology.

**Figure S11**. BioGeoBEARS BAYAREALIKE+J on Asparagaceae maximum likelihood chloroplast gene (Ml cpDNA) supermatrix.

**Table S1**. Sampling information, source, voucher, etc. for each sample used in the analysis. Note: we have not added the information for the samples included that were sequenced in previous studies.

**Table S2**. Summary statistics for nuclear and chloroplast datasets. Information includes recovery rates for each sample as well as sequence length of each locus for each sample.

**Table S3**. Divergence times and biogeographic reconstruction probabilities for each subfamily across each topology (ML nuclear, ML cpDNA, wASTRAL).

**Table S4**. Biogeographic model comparison for each topology.

