## Supplementary material for "Panic at the Discordance: the Asparagaceae Phylogenetic Backbone": Figure S9

**BioGeoBEARS BAYAREALIKE+J on Asparagaceae wASTRAL Topology**
**ancstates: global optim, 6 areas max. d=0.004; e=0.0047; j=0.0108; LnL=−266.23**

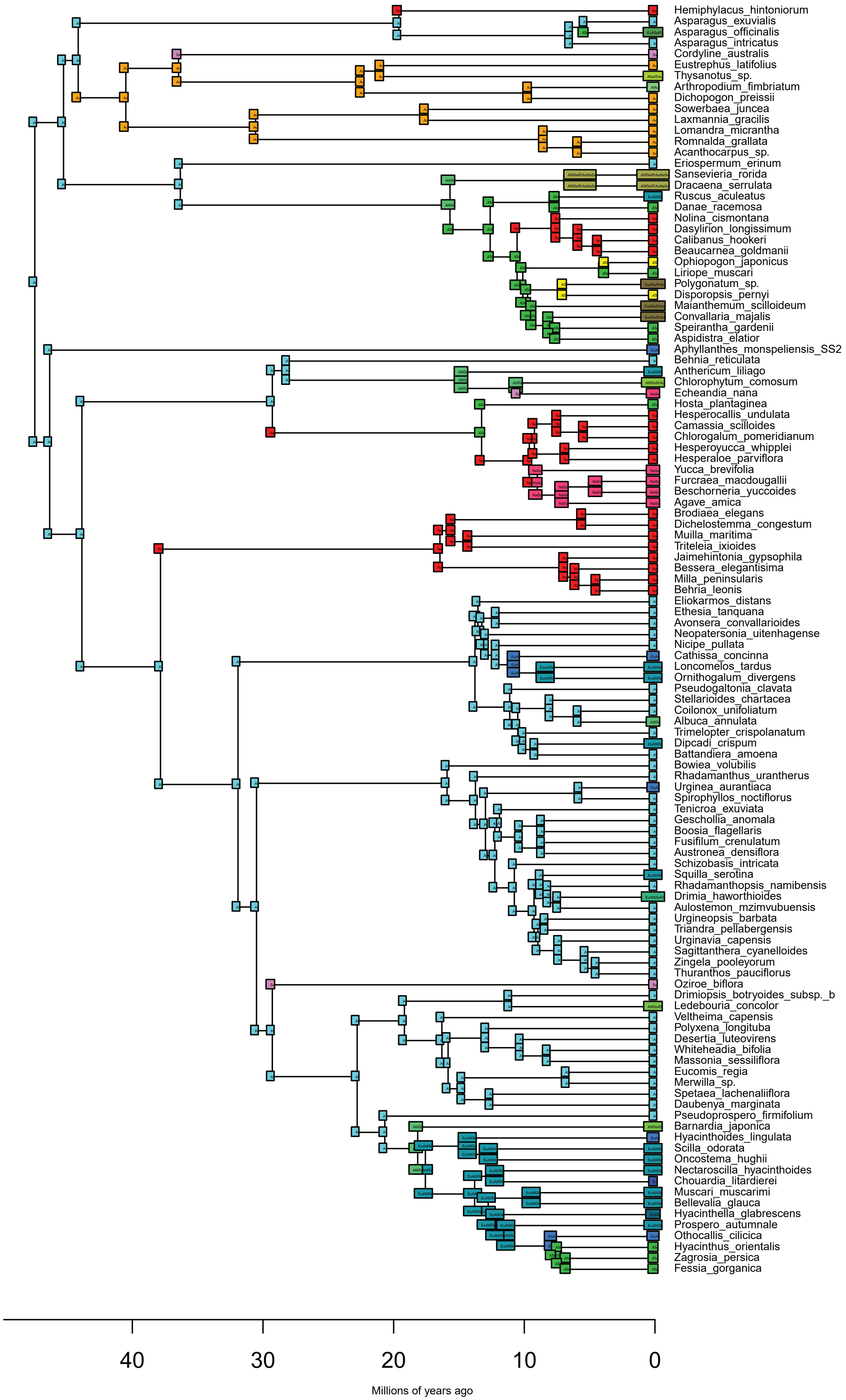
