## Supplementary material for "Panic at the Discordance: the Asparagaceae Phylogenetic Backbone": Figure S11

**BioGeoBEARS BAYAREALIKE+J on Asparagaceae ML cpDNA Topology**
**ancstates: global optim, 6 areas max. d=0.0031; e=0.0042; j=0.0119; LnL=−276.63**

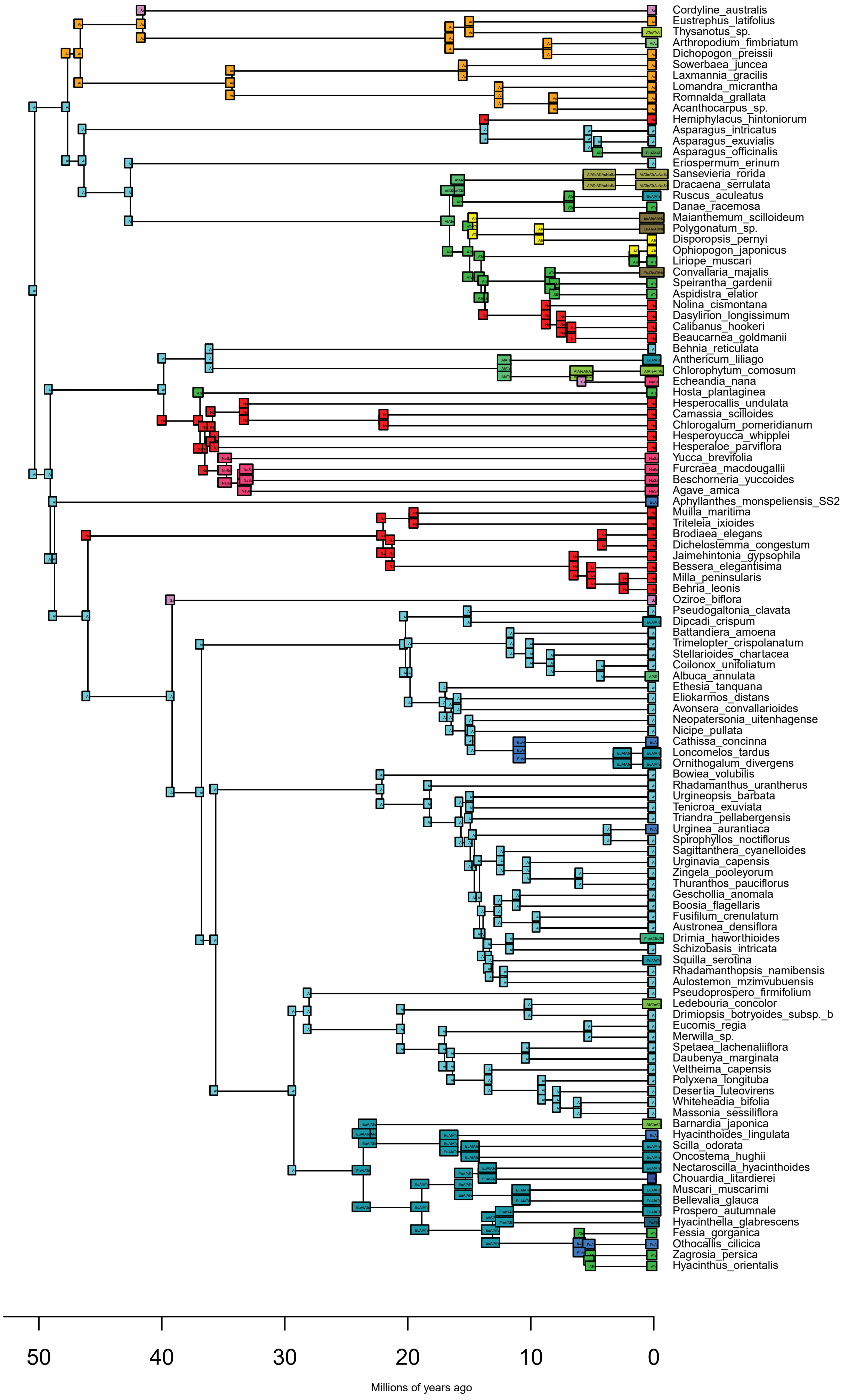
