## Supplementary material for "Panic at the Discordance: the Asparagaceae Phylogenetic Backbone": Table S1

| code | genus | species | species_name | taxonomy | supplier | leaf type | collection_info | misc_notes |
|---|---|---|---|---|---|---|---|---|
| MMAR | Muilla | maritima | Muilla_maritima | Brodiaeoideae | Harvey | silica-leaf | CCH s.n. | |
| CHOO | Calibanus | hookeri | Calibanus_hookeri | Nolinoideae | Harvey | silica-leaf | CCH s.n. | |
| SROR | Sansevieria | rorida | Sansevieria_rorida | Nolinoideae | Harvey | silica-leaf | Lavranos 23319; Somalia | |
| BGOL | Beaucarnea | goldmanii | Beaucarnea_goldmanii | Nolinoideae | Harvey | silica-leaf | CCH s.n. | |
| SODO | Scilla | odorata | Scilla_odorata | Scilloideae | Harvey | silica-leaf | CCH s.n. | |
| DSER | Dracaena | serrulata | Dracaena_serrulata | Nolinoideae | Harvey | silica-leaf | ISI 2004-20 | |
| EERI | Eriospermum | erinum | Eriospermum_erinum | Nolinoideae | Harvey | silica-leaf | cultivated | |
| HUND | Hesperocallis | undulata | Hesperocallis_undulata | Agavoideae | Sierra Jaeger | silica-leaf | Sierra Jaeger 26 | Painted Canyon Rd, CA: 13 miles from Box Canyon Rd |
| HPLA | Hosta | plantaginea | Hosta_plantaginea | Agavoideae | MacNeill | silica-leaf | UNA00075352 | Collected from MOBOT |
| PZAP | Agave | zapopanensis | Agave_zapopanensis | Agavoideae | MacNeill | silica-leaf | PZAPA1-6585-63 | |
| PRSPE | Prochnyanthes | species | Prochnyanthes_species | Agavoideae | MacNeill | silica-leaf | PROCNOVA1-7588AR-68 | |
| ENAN | Echeandia | nana | Echeandia_nana | Agavoideae | MacNeill | silica-leaf | Aarón Rodriguez et al. 6709, IBUG 205048 | |
| EECH | Echeandia | echinoides | Echeandia_echinoides | Agavoideae | MacNeill | silica-leaf | Aarón Rodriguez & L. Ortiz-Catedral 3019, IBUG 156725 | |
| MJAL | Agave | jaliscana | Agave_jaliscana | Agavoideae | MacNeill | silica-leaf | 4835-13 | |
| EDIS | Eliokarmos | distans | Eliokarmos_distans | Scilloideae | Martinez-Azori | silica-leaf | M422 | |
| NPUL | Nicipe | pullata | Nicipe_pullata | Scilloideae | Martinez-Azori | silica-leaf | M1306 | |
| GANO | Geschollia | anomala | Geschollia_anomala | Scilloideae | Martinez-Azori | silica-leaf | M1343 | |
| BFLA | Boosia | flagellaris | Boosia_flagellaris | Scilloideae | Martinez-Azori | silica-leaf | M1459 | |
| UBAR | Urgineopsis | barbata | Urgineopsis_barbata | Scilloideae | Martinez-Azori | silica-leaf | M1659 | |
| CUNI | Coilonox | unifoliatum | Coilonox_unifoliatum | Scilloideae | Martinez-Azori | silica-leaf | M1885 | |
| ETAN | Ethesia | tanquana | Ethesia_tanquana | Scilloideae | Martinez-Azori | silica-leaf | M815 | |
| TEXU | Tenicroa | exuviata | Tenicroa_exuviata | Scilloideae | Martinez-Azori | silica-leaf | M1913 | |
| ZPOO | Zingela | pooleyorum | Zingela_pooleyorum | Scilloideae | Martinez-Azori | silica-leaf | M1710 | |
| WBIF | Whiteheadia | bifolia | Whiteheadia_bifolia | Scilloideae | Martinez-Azori | silica-leaf | M1510 | |
| BAMO | Battandiera | amoena | Battandiera_amoena | Scilloideae | Martinez-Azori | silica-leaf | MA s.n. | |
| RURA | Rhadamanthus | urantherus | Rhadamanthus_urantherus | Scilloideae | Martinez-Azori | silica-leaf | M892 | |
| RNAM | Rhadamanthops | namibensis | Rhadamanthopsis_namibensis | Scilloideae | Martinez-Azori | silica-leaf | M1593 | |
| DCRI | Dipcadi | crispum | Dipcadi_crispum | Scilloideae | Martinez-Azori | silica-leaf | M647 | |
| TPEL | Triandra | pellabergensis | Triandra_pellabergensis | Scilloideae | Martinez-Azori | silica-leaf | M1671 | |
| HLIN | Hyacinthoides | lingulata | Hyacinthoides_lingulata | Scilloideae | Martinez-Azori | silica-leaf | M2 | |
| SNOC | Spirophyllos | noctiflorus | Spirophyllos_noctiflorus | Scilloideae | Martinez-Azori | silica-leaf | AG4 | |
| SCYA | Sagittanthera | cyanelloides | Sagittanthera_cyanelloides | Scilloideae | Martinez-Azori | silica-leaf | MA s.n. | van Jaarsveld |
| AMZI | Aulostemon | mzimbuvuensis | Aulostemon_mzimvubuensis | Scilloideae | Martinez-Azori | silica-leaf | MA s.n. | TYPE van Jaarsveld |
| MSES | Massonia | sessiliflora | Massonia_sessiliflora | Scilloideae | Martinez-Azori | silica-leaf | M1860 | |
| UCAP | Urginavia | capensis | Urginavia_capensis | Scilloideae | Martinez-Azori | silica-leaf | M1332 | |
| POBT | Prospero | obtusifolium | Prospero_obtusifolium | Scilloideae | Martinez-Azori | silica-leaf | M1949 | |
| FCRE | Fusifilum | crenulatum | Fusifilum_crenulatum | Scilloideae | Martinez-Azori | silica-leaf | M1568 | |
| UAUR | Urginea | aurantiaca | Urginea_aurantiaca | Scilloideae | Martinez-Azori | silica-leaf | M1988 | |
| PCLA | Pseudogaltonia | clavata | Pseudogaltonia_clavata | Scilloideae | Martinez-Azori | silica-leaf | W1530 | |
| OBIF | Oziroe | biflora | Oziroe_biflora | Scilloideae | Martinez-Azori | silica-leaf | W3471 | |
| AANN | Albuca | annulata | Albuca_annulata | Scilloideae | Martinez-Azori | silica-leaf | M241 | |
| DLUT | Desertia | luteovirens | Desertia_luteovirens | Scilloideae | Martinez-Azori | silica-leaf | M1591 | |
| DHAW | Drimia | haworthioides | Drimia_haworthioides | Scilloideae | Martinez-Azori | silica-leaf | M978 | |
| LTAR | Loncomelos | tardus | Loncomelos_tardus | Scilloideae | Martinez-Azori | silica-leaf | W5078 | |
| ODIV | Ornithogalum | divergens | Ornithogalum_divergens | Scilloideae | Martinez-Azori | silica-leaf | MA s.n. | |
| PFIR | Pseudoprosperc | firmifolium | Pseudoprospero_firmifolium | Scilloideae | Martinez-Azori | silica-leaf | W4368 | |
| SCHA | Stellarioides | chartacea | Stellarioides_chartacea | Scilloideae | Martinez-Azori | silica-leaf | M990 | |
| MSPE | Merwilla | species | Merwilla_species | Scilloideae | Martinez-Azori | silica-leaf | M1354 | |
| NUIT | Neopatersonia | uitenhagense | Neopatersonia_uitenhagense | Scilloideae | Martinez-Azori | silica-leaf | W3887 | |
| TCRI | Trimelopter | crispolanatum | Trimelopter_crispolanatum | Scilloideae | Martinez-Azori | silica-leaf | M1781 | |
| UFUG | Urginea | fugax | Urginea_fugax | Scilloideae | Martinez-Azori | silica-leaf | M1938 | |
| ADEN | Austronea | densiflora | Austronea_densiflora | Scilloideae | Martinez-Azori | silica-leaf | M945 | |
| TPAU | Thuranthos | pauciflorus | Thuranthos_pauciflorus | Scilloideae | Martinez-Azori | silica-leaf | W5210 | |

| | | | | | | | | |
|---|---|---|---|---|---|---|---|---|
| BGAR | Bowiea | gariepensis | Bowiea_gariepensis | Scilloideae | Martinez-Azori | silica-leaf | M1673 | |
| SSER | Squilla | serotina | Squilla_serotina | Scilloideae | Martinez-Azori | silica-leaf | M1941 | |
| DLON | Dasylirion | longissimum | Dasylirion_longissimum | Nolinoideae | Könyves | silica-leaf | W20061423-A | |
| DRAC | Danae | racemosa | Danae_racemosa | Nolinoideae | Könyves | silica-leaf | JCD s.n. | |
| CMAJ | Convallaria | majalis | Convallaria_majalis | Nolinoideae | Könyves | silica-leaf | JCD s.n. | |
| ALIL | Anthericum | liliago | Anthericum_liliago | Agavoideae | Könyves | silica-leaf | Major' | |
| ASPE | Allium | peninsulare | Allium_peninsulare | Amaryllidaceae | Könyves | silica-leaf | W20070037-A | |
| BCAL | Triteleia | ixioides | Triteleia_ixioides | Brodiaeoideae | Könyves | silica-leaf | W20060744-A | |
| TIXI | Brodiaea | elegans | Brodiaea_elegans | Brodiaeoideae | Könyves | silica-leaf | W19990684-A | |
| ZPER | Zagrosia | persica | Zagrosia_persica | Scilloideae | Könyves | silica-leaf | W940226-A | |
| PAUT | Prospero | autumnale | Prospero_autumnale | Scilloideae | Könyves | silica-leaf | W953134-A | |
| OCIL | Othocallis | cilicica | Othocallis_cilicica | Scilloideae | Könyves | silica-leaf | W200073120-A | |
| OHUG | Oncostema | hughii | Oncostema_hughii | Scilloideae | Könyves | silica-leaf | JCD s.n. | |
| NHYA | Nectaroscilla | hyacinthoides | Nectaroscilla_hyacinthoides | Scilloideae | Könyves | silica-leaf | JCD s.n. | |
| HGLA | Hyacinthella | glabrescens | Hyacinthella_glabrescens | Scilloideae | Könyves | silica-leaf | W20003783-A | |
| HORI | Hyacinthus | orientalis | Hyacinthus_orientalis | Scilloideae | Könyves | silica-leaf | W20122095-A | |
| FGOR | Fessia | gorganica | Fessia_gorganica | Scilloideae | Könyves | silica-leaf | W20060782-A | |
| CLIT | Chouardia | litardierei | Chouardia_litardierei | Scilloideae | Könyves | silica-leaf | W901608-A | |
| BJAP | Barnardia | japonica | Barnardia_japonica | Scilloideae | Könyves | silica-leaf | W20081977-A | |
| SGAR | Speirantha | gardenii | Speirantha_gardenii | Nolinoideae | Könyves | silica-leaf | JCD s.n. | |
| DPER | Disporopsis | pernyi | Disporopsis_pernyi | Nolinoideae | Könyves | silica-leaf | W20190744-A | |
| RACU | Ruscus | aculeatus | Ruscus_aculeatus | Nolinoideae | Könyves | silica-leaf | Sparkler = W943656-C | |
| POSPE | Polygonatum | species | Polygonatum_species | Nolinoideae | Howard | silica-leaf | CCH s.n. | |
| LMIC | Lomandra | micrantha | Lomandra_micrantha | Lomandroideae | Birch | silica-leaf | JLB921 (MELU155159a) | |
| CSCI | Camassia | scilloides | Camassia_scilloides | Agavoideae | Howard | silica-leaf | CCH s.n. | |
| SJUN | Sowerbaea | juncea | Sowerbaea_juncea | Lomandroideae | Birch | silica-leaf | JLB952 (MELU155162a) | |
| LGRA | Laxmannia | gracilis | Laxmannia_gracilis | Lomandroideae | Birch | silica-leaf | JLB944 (MELU155161a) | |
| RGRA | Romnalda | grallata | Romnalda_grallata | Lomandroideae | Birch | silica-leaf | BF Gunn 1268 (MEL2494959) | |
| CAUS | Cordyline | australis | Cordyline_australis | Lomandroideae | Birch | silica-leaf | JLB892 (MELU155158a) | |
| AFIM | Arthropodium | fimbriatum | Arthropodium_fimbriatum | Lomandroideae | Birch | silica-leaf | JLB942 (MELU155160a) | |
| JGYP | Jaimehintonia | gypsophila | Jaimehintonia_gypsophila | Brodiaeoideae | Gándara | silica-leaf | Ortiz-Brunel 1695 | |
| MPEN | Milla | peninsularis | Milla_peninsularis | Brodiaeoideae | Gándara | silica-leaf | E. Gándara 3311 | |
| MSCI | Maianthemum | scilloides | Maianthemum_scilloides | Nolinoideae | Gándara | silica-leaf | A. Morales-González 3 | |
| BELE | Bessera | elegantisima | Bessera_elegantisima | Brodiaeoideae | Gándara | silica-leaf | E. Ruiz-Sanchez 654 | |
| BTUI | Bessera | tuitensis | Bessera_tuitensis | Brodiaeoideae | Gándara | silica-leaf | E. Gándara 3282 | |
| BLEO | Behria | leonis | Behria_leonis | Brodiaeoideae | Gándara | silica-leaf | E. Gándara 3300 | |
| BTEN | Behria | tenuiflora | Behria_tenuiflora | Brodiaeoideae | Gándara | silica-leaf | E. Gándara 3306 | |
| MMUS | Muscari | muscarimi | Muscari_muscarimi | Scilloideae | Könyves | silica-leaf | W918287-A | |
| BGLA | Bellevalia | glauca | Bellevalia_glauca | Scilloideae | Könyves | silica-leaf | W20030942-A | |
| YBRE | Yucca | brevifolia | Yucca_brevifolia | Agavoideae | California Bota | silica-leaf | Calbg 23873 A20 | |
| NCIS | Nolina | cismontana | Nolina_cismontana | Nolinoideae | California Bota | silica-leaf | Calbg 17833*A2 | |
| HPEN | Hesperoyuuca | peninsularis | Hesperoyuuca_peninsularis | Agavoideae | California Bota | silica-leaf | Calbg 24707 | |
| 4450 | Avonsera | convallarioides | Avonsera_convallarioides | Scilloideae | Martinez-Azori | DNA | 4450 | suspended in TE |
| G1 | Cathissa | concinna | Cathissa_concinna | Scilloideae | Martinez-Azori | DNA | G1 | suspended in TE |
| COBL | Cyrtanthus | obliquus | Cyrtanthus_obliquus | Amaryllidaceae | Howard | silica-leaf | CCH s.n. | ex Tim Harvey; cultivated |
| 310 | Daubenya | marginata | Daubenya_marginata | Scilloideae | Martinez-Azori | DNA | 310 | suspended in TE |
| W3948 | Eucomis | regia | Eucomis_regia | Scilloideae | Martinez-Azori | DNA | W3948 | suspended in TE |
| K2 | Ornithogalum | baeticum | Ornithogalum_baeticum | Scilloideae | Martinez-Azori | DNA | K2 | suspended in TE |
| W4327 | Polyxena | longituba | Polyxena_longituba | Scilloideae | Martinez-Azori | DNA | W4327 | suspended in TE |
| W2512 | Rhodocodon | linearifolius | Rhodocodon_linearifolius | Scilloideae | Martinez-Azori | DNA | W2512 | suspended in TE |
| 4903 | Spetaea | lachenaliifolia | Spetaea_lachenaliiflora | Scilloideae | Martinez-Azori | DNA | 4903 | suspended in TE |
| 2341 | Veltheima | capensis | Veltheima_capensis | Scilloideae | Martinez-Azori | DNA | 2341 | suspended in TE |
| ZCAN | Zephyranthes | candida | Zephyranthes_candida | Amaryllidaceae | Howard | silica-leaf | CCH s.n. | cultivated material from yard |
| | Agave | amica | Agave_amica | Agavoideae | Bentz | DNA | | SRA: PRJNA1292917 |

| | | | | | | | |
|---|---|---|---|---|---|---|---|
| Agave | moranii | Agave_moranii | Agavoideae | Salywon | DNA | DBG 1991-0471-0202 | SRA: PRJNA1292931 |
| Agave | scabra | Agave_scabra | Agavoideae | Bentz | DNA | | SRA: PRJNA1292918 |
| Agave | xylocantha | Agave_xylocantha | Agavoideae | Salywon | DNA | DBG 1962-7168-02-07 | SRA: PRJNA1292935 |
| Furcraea | hexapetala | Furcraea_hexapetala | Agavoideae | Salywon | DNA | DBG 1987-0847-02-25 | SRA: PRJNA1423816 |
| Furcraea | macdougallii | Furcraea_macdougallii | Agavoideae | Bentz | DNA | | SRA: PRJNA1423819 |
| Hesperaloe | funifera | Hesperaloe_funifera | Agavoideae | Salywon | DNA | DBG 2019-0695-01 | SRA: PRJNA1423817 |
| Hesperaloe | parviflora | Hesperaloe_parviflora | Agavoideae | Salywon | DNA | DBG 1974-015401401 | SRA: PRJNA1423815 |
| Agave | virginica | Agave_virginica | Agavoideae | Bentz | | P.C. Bentz 126 (Bentz and Leebens-Mack, 2024) | |
| Agave | xylocantha | Agave_xylocantha | Agavoideae | Publicly available | | | |
| Agave | gracielae | Agave_gracielae | Agavoideae | Publicly available | | | |
| Agave | virginica | Agave virginica | Agavoideae | Publicly available | | | |
| Behnia | reticulata | Behnia_reticulata | Agavoideae | Bentz | | Burrows & Burrows 15952 (Bentz and Leebens-Mack, 2024) | |
| Beschorneria | yuccioides | Beschorneria_yuccioides | Agavoideae | Bentz | | K. Heyduk 3 | |
| Beschorneria | septentrionalis | Beschorneria_septentrionalis | Agavoideae | (Bentz) JLM freezer | | JLM s.n. | |
| Chlorogalum | pomeridianum | Chlorogalum_pomeridianum | Agavoideae | (Bentz) JLM freezer | | JLM s.n. | |
| Chlorophytum | comosum | Chlorophytum_comosum | Agavoideae | Bentz | | P.C. Bentz 127 (Bentz and Leebens-Mack, 2024) | |
| Hesperoyucca | whipplei | Hesperoyucca_whipplei | Agavoideae | Bentz | JLM freezer | JLM s.n. | |
| Yucca | filamentosa | Yucca_filamentosa | Agavoideae | Bentz | | P.C. Bentz 132 (Bentz and Leebe | Note: In Bentz & Leebens-Mack 2024, collection number is P.C. Bentz 131; corrected here to PCB 132. |
| Yucca | aloifolia | Yucca_aloifolia | Agavoideae | Bentz | | P.C. Bentz 131 | |
| Hemiphylacus | hintoniorum | Hemiphylacus_hintoniorum | Asparagoideae | Bentz | | P.C. Bentz 55 | |
| Asparagus | officinalis | Asparagus_officinalis | Asparagoideae | Bentz | | P.C. Bentz 81 (Bentz and Leebens-Mack, 2024) | |
| Asparagus | bayeri | Asparagus_bayeri | Asparagoideae | Bentz | | Burrows, J.E. & S.M. 11919 (Bentz and Leebens-Mack, 2024) | |
| Asparagus | concinnus | Asparagus_concinnus | Asparagoideae | Bentz | | Burrows, J.E. & S.M. 15358 (Bentz and Leebens-Mack, 2024) | |
| Asparagus | exuvialis | Asparagus_exuvialis | Asparagoideae | Bentz | | Burrows, J.E. & S.M. 15913 (Bentz and Leebens-Mack, 2024) | |
| Dichelostemma | congestum | Dichelostemma_congestum | Brodiaeoideae | Bentz | | D. Hee 08 (Bentz and Leebens-Mack, 2024) | |
| Triteleia | laxa | Triteleia_laxa | Brodiaeoideae | Bentz | | R. Halse 8737 (Bentz and Leebens-Mack, 2024) | |
| Acanthocarpus | sp | Acanthocarpus_sp | Lomandroideae | Bentz | JLM freezer | K. Thiele s.n. (Bentz and Leebens-Mack, 2024) | |
| Eustrephus OR | latifolius | Eustrephus OR Chamaescilla_latifolius | Lomandroideae | Bentz | | W. Zomlefer 2326 (Bentz and Leebens-Mack, 2024) | |
| Thysanotus | sp | Thysanotus_sp | Lomandroideae | Bentz | JLM freezer | K. Thiele s.n. (Bentz and Leebens-Mack, 2024) | |
| Dichopogon | preissii | Dichopogon_preissii | Lomandroideae | Bentz | JLM freezer | K. Thiele RRT3701 (Bentz and Leebens-Mack, 2024) | |
| Liriope | muscari | Liriope_muscari | Nolinoideae | Bentz | | P.C. Bentz 129 (Bentz and Leebens-Mack, 2024) | |
| Aspidistra | elatior | Aspidistra_elatior | Nolinoideae | Bentz | | P.C. Bentz 130 (Bentz and Leebens-Mack, 2024) | |
| Ophiopogon | japonicus | Ophiopogon_japonicus | Nolinoideae | Bentz | | P.C. Bentz 128 (Bentz and Leebens-Mack, 2024) | |
| Albuca | cf. recurva | Albuca_cf. recurva | Scilloideae | Howard | silica-leaf | C.C. Howard 205 (Bentz and Leebens-Mack, 2024) | |
| Bowiea | volubilis | Bowiea_volubilis | Scilloideae | Howard | silica-leaf | C.C. Howard 156 (Bentz and Leebens-Mack, 2024) | |
| Dipcadi | sp. | Dipcadi_sp. | Scilloideae | Howard | silica-leaf | C.C. Howard 222 (Bentz and Leebens-Mack, 2024) | |
| Drimia | intricata | Drimia_intricata | Scilloideae | Howard | silica-leaf | C.C. Howard HBG90233 (Bentz and Leebens-Mack, 2024) | |
| Drimiopsis | botryoides subs | Drimiopsis_botryoides subsp. botryoide | Scilloideae | Howard | silica-leaf | C.C. Howard 154 (Bentz and Leebens-Mack, 2024) | |
| Ledebouria | nitida | Ledebouria_nitida | Scilloideae | Howard | silica-leaf | C.C. Howard 246 | |
| Ledebouria | scabrida | Ledebouria_scabrida | Scilloideae | Howard | silica-leaf | C.C. Howard s.n. (Bentz and Leebens-Mack, 2024) | |
| Schizobasis | intricata | Schizobasis_intricata | Scilloideae | Howard | silica-leaf | Huntington Botanical Garden | |
| Dianella | tasmanica | Dianella_tasmanica | Asphodelaceae | Bentz | JLM freezer | T. Givnish s.n. (Bentz and Leebens-Mack, 2024) | |
| Aphyllanthes | monspeliensis | Aphyllanthes_monspeliensis | Aphyllanthoide | Martinez-Azori | silica-leaf | M. Martinez-Azorin RB1 | |
| Aphyllanthes | monspeliensis | Aphyllanthes_monspeliensis | Aphyllanthoide | Martinez-Azori | silica-leaf | M. Martinez-Azorin RB2 | |
| Aphyllanthes | monspeliensis | Aphyllanthes_monspeliensis | Aphyllanthoide | Martinez-Azori | silica-leaf | M. Martinez-Azorin RB3 | |
| Aphyllanthes | monspeliensis | Aphyllanthes_monspeliensis | Aphyllanthoide | Martinez-Azori | silica-leaf | M. Martinez-Azorin SS1 | |
| Aphyllanthes | monspeliensis | Aphyllanthes_monspeliensis | Aphyllanthoide | Martinez-Azori | silica-leaf | M. Martinez-Azorin SS2 | |
| Aphyllanthes | monspeliensis | Aphyllanthes_monspeliensis | Aphyllanthoide | Martinez-Azori | silica-leaf | M. Martinez-Azorin SS3 | |
